# Detection and quantification of the heterogeneity of *S. aureus* bacterial populations to identify antibiotic-induced persistence

**DOI:** 10.1101/320093

**Authors:** Marwa M. Hassan, Mark S. Butler, Andrea Ranzoni, Matthew A. Cooper

## Abstract

**Objectives:** Persister cells are characterised as being viable but non-culturable, a state that preserves their metabolic energy to survive the environmental stress, which allows for recurrent infections. Detection of persisters is, therefore, not possible with standard culture-dependent methods. Furthermore, the effect of antibiotics on the induction of persisters has not been assessed. This study aimed to identify antibiotic-induced persistence and determine the percentage of heterogeneity.

**Methods:** Vancomycin, daptomycin and dalbavancin were assessed by standard MIC methods against selected *Staphylococcus aureus* strains. Replicates of MIC assays were stained with propidium iodide to quantify live/dead and a reactive oxygen species (ROS) dye to detect and quantify persisters using culture-independent single-cell sorting, independently. A comparative analysis was then performed.

**Results:** Dalbavancin showed the lowest MIC values against tested *S. aureus* strains followed by daptomycin and vancomycin. Cell sorting of vancomycin-, daptomycin- and dalbavancin-treated *S. aureus* strains showed a range of 1.9–10.2%, 17.7–62.9% and 7.5–77.6% live cells based on the strain, respectively, in which daptomycin, in particular, was a strong inducer of a persister population. Persisters represented 3.7–16% of the bacterial population.

**Conclusions:** The culture-independent identification of antibiotic-induced persistence through studying at the single-cell level showed different efficacy of antibiotics than standard MIC. Vancomycin was the most effective antibiotic against tested strains followed by dalbavancin then daptomycin as assessed by cell sorting. Therefore, re-evaluation of standard MIC methods may be required to assess the efficacy of antibiotics. Additionally, the detection of daptomycin-associated persisters may provide an elucidation to the reported rapid resistance development *in vivo*.

## Introduction

Persistence is defined as the ability of a genetically identical sub-population of the bacterial cells to survive the stress of the antibiotics and die at a slower rate than the rest of the population.^1^ In time-kill experiments, a sharp decrease in the percentage of live cells occurs followed by a plateau survival rate due to the slow rate of death of these persister cells. Tolerance, however, is the transient ability of all bacterial populations to adapt to the environmental pressure or antibiotics stress and remain alive, which can result in the acquisition of new mutations. Prolonged exposure to the same stress diminishes this survival ability.^2^ The distinction between persistence and tolerance has been made by the Balaban group, in which persistence was proposed to be a higher level of tolerance with a lower percentage of viable cells during the environmental stress.^2^ Accordingly, the persistent/tolerant live bacterial population can adapt to antibiotics at a higher concentration than the minimum inhibitory concentration (MIC), while maintaining a sensitive antibiotic profile.^2^ Thus, Brauner *et al*. proposed a model to define tolerance and persistence based on the duration required to kill 99% and 99.99% of the bacterial population.^2^ This 0.01–1% of the bacterial population can switch from non-culturable to a culturable state once the environmental stress has been eliminated.^2^

Persisters have been characterised by their reduced metabolism and growth rate, in order to save energy.^3^ They were also shown to stop their replication and become viable but non-culturable,^4^ a dormant stage that has been associated with biofilm formation.^5,6^ Both tolerance and persistence have been linked to antibiotic resistance and recurrent infections.^7,8^ Persistence has also been associated with the formation of a small colony variant (SCV), which is characterised by a slow growth rate.^9^ SCV strains are also associated with recurrent infections,^10,11^ due to their decreased electron transport activity, lower respiration rate and the ability to return to the parent phenotype.^9–11^

The phenomena of persistence is not new, it is an inherent ability of some bacterial species, such as *Mycobacterium tuberculosis*;^12^ however, persistence has also been shown in other bacterial species such as *Escherichia coli*,^3^ *S. aureus*,^13^ *Clostridium difficile* and *Salmonella* species.^12,14,15^ The molecular mechanisms underlying persistence are now being extensively studied on these rapid growing model species. The survival of these persisters is based on few shared key molecular mechanisms, which are based on oxidative stress, toxin-antitoxin (TA) modules and quorum sensing.^4,5,16,17^ The oxidative stress is a well characterised mechanism, especially with antibiotics. Under antibiotic stress, cytotoxic ROS are generated to target bacterial proteins, lipids and DNA, which lead to cell damage and subsequently death.^16^ However, persisters have the ability to suppress the oxidative stress pathway through either scavenging the generated ROS or producing less toxic radicals.^16^ Lower levels of oxidative stress and free radicals prevents DNA damage, which allows the cells to survive the bactericidal effect of antibiotics. This mechanism has been reported in *S. aureus*, *E. coli* and *Pseudomonas aeruginosa*.^16^

The detection of these non-culturable cells is not possible by culture-based methods, and their estimated low percentage, 0.01–1%, necessitated their detection using single-cell approaches. Recent studies have developed single-cell microfluidics, microscopy and cell sorting methods using fluorescence-activated cell sorting (FACS) to detect and characterise persister cells.^3,18–20^ FACS has also been used to analyse the metabolic and growth state of persisters,^21^ identify heterogeneity in cell division, which protected the cells from the killing effect of the complement system and antibiotics,^22^ and align cell sorting with sequencing to study gene expression of persisters.^23^ The application of single-cell techniques to detect persistence has been reviewed elsewhere.^24,25^ Recently, FACS was applied to detect carbapenem resistance in *Klebsiella pneumoniae*.^26^

This study aimed to re-assess the standard MIC methods and investigate the role of three cell-wall antibiotics in the formation of persistence against *S. aureus* strains. A culture-independent cell sorting method was developed to assess, detect and quantify live, dead and persister cells in MIC assays. This quantification data was then correlated with the MIC values of these antibiotics by the broth micro- and macrodilution methods. The results showed a single-cell based method that could be used for an in-depth evaluation of the efficacy of antibiotics, based on both the percentage of live/dead cells and persistence formation. The single-cell sorting of MIC assays showed that the standard MIC methods may not reflect the potency of antibiotics, show bacterial heterogeneity and/or detect antibiotic associated persistence.

## Methods

### 1- Determination of MIC using standard methods

Determination of MIC was performed using the broth micro- and macrodilution methods according to the Clinical and Laboratory Standards Institute (CLSI) guidlines.^27^ All daptomycin assays were supplemented with calcium to a final concentration of 50 mg/L.^28^ Sensitive and resistant *S. aureus* strains used in this study are listed in **Table S1**.

To determine the MIC by resazurin, a final concentration of 0.04 mM (0.01 mg/mL) resazurin (Sigma Aldrich) was added aseptically to all MIC plates and tubes, and further incubated at 37 ºC in the dark. All MIC plates and tubes were checked for colour change at 4 and 24 h. All tests were repeated with two independent cultures and each tested in duplicate.

### 2- Detection of MIC using macrodilution method by FACS

The bacterial cells of the broth macrodilution MIC replicates were washed twice from media by centrifugation at a maximum speed for 10 min at room temperature and resuspended in 200 µL of sterile 5 mM HEPES buffer containing 20 mM glucose, after the 24 h incubation. A subculture of an overnight bacterial culture was washed as previously described as used as a positive control of live cells. Then, an aliquote of the bacterial subculture was centrifuged and treated with 80% ethanol for 2 h before washed as previously described and used as the control dead cells. Then, samples were analysed for live/dead by staining with propidium iodide (PI) (Sigma) dye at a final concentration of 8 mg/L and incubated for 5 min. Samples were detected on BD FACSCanto II flow cytometer (BD Biosciences) and analysed using Kaluza Analysis 1.3 software (Beckman Coulter).

Unstained bacterial cells were compared using the FSC and SSC (forward and side scatter) gating (**Figure S1a**). Gate ‘C’ represents live cells and gate ‘D’ represents dead cells (**Figure S1b**). Negative control for media and buffer were used to confirm that the signal is due to the presence of bacteria. The gates were slightly modified to compensate for the difference between the antibiotics killing mode of action and a maximum of 10K events was used to standardise the number of detected events.

### 3- Detection of antibiotic stress level and persistence by FACS

The bacterial cells of the broth macrodilution MIC replicates were washed as previously described and stained with CELLROX green dye (Life Technology). CELLROX dye was added at a final concentration of 2.5 µM and incubated for 5 min before sorted. A maximum of 10K events was used to standardise the number of detected events. Unstained bacterial cells were compared using the FSC and SSC gating.

## Results and discussions

The main aim of this study was to identify antibiotic-induced persistence through evaluating the current standard MIC methods and assessing the efficacy of antibiotics independent of the culture methods. Vancomycin, a bacteriostatic^29^ or slow bactericidal^30^ glycopeptide, daptomycin, a bactericidal lipopeptide^31,32^ and dalbavancin, a bactericidal lipoglycopeptide,^33,34^ were chosen for this study due to their bacterial membrane targeting and variable efficacy against selected *S. aureus* strains.

### 1. MIC determination by standard methods

The determination of the MIC was performed by the two-standard broth microdilution and macrodilution methods to detect and demonstrate the difference in MIC values. Since, the standard endpoint reading of both methods is by visual identification of the lowest antibiotic concentration that shows no growth, resazurin dye was added aseptically to assess the sensitivity of visual detection. Resazurin is a redox dye that changes its colour upon reduction by live cells and was used in MIC determination but not considered as a standard protocol.^35,36^ The final concentration of the dye was optimised for the lowest detection limit through serial dilutions of resazurin and bacterial cells. The results showed that 40 µM resazurin (as a final concentration) had the lowest possible detection limit, which was 10^7^ cfu/mL live cells and a higher concentration of resazurin hinders the detectable colour change.

Vancomycin MICs by resazurin broth macrodilution showed a 2-fold increase in MIC values with MRSA and hVISA compared to the microdilution method, differentiating the hVISA strain from MSSA strains due to the higher number of live cells (**Table 1**). The phenotypic distinction between some of the hVISA and MSSA strains has always been difficult due to their identical sensitive MIC rather to an intermediate cut-off using the broth microdilution method.^37,38^ However, these results require an extensive study with a larger number of heterogeneous strains to prove distinction.

Daptomycin showed an increased MIC values of 2- to 4-fold as measured by the resazurin broth macrodilution method (**Table 1**). All tested strains had MIC values of 1 mg/L while hVISA and VISA strains had MIC of 4 and 8 mg/L, respectively (**Table 1**). These MIC results suggest that both strains may have a daptomycin intermediate or resistance profile; however, ≤ 1 mg/L is the only reported^27^ daptomycin-sensitive breakpoint.

Dalbavancin MIC values increased by 4-fold against all the strains using the resazurin macrodilution method. Even though the detailed breakpoint of dalbavancin’s intermediate-resistance has not been identified, MSSA and MRSA showed MIC value of 0.25 mg/L, whereas hVISA and VISA strains had MIC of 0.5 and 1 mg/L, respectively. VRSA strains showed MIC of 4 and 8 mg/L (**Table 1**). The increase in dalbavancin MICs with increased vancomycin resistance is due to the similarity in their mode of action in binding to the D-alanyl-D-alanine residues of the bacterial cell wall.^39,40^ However, dalbavancin anchors and dimerises in the bacterial membrane, which results in a higher binding affinity and effectiveness against VRSA.^39^

These results showed higher MIC values for the broth macrodilution compared to the microdilution method (**Table 1**), as previously reported.^41,42^ Vancomycin, daptomycin and dalbavancin had an increased MIC of 2-, 2–4 and 4-fold with the resazurin macrodilution method. The increased MIC-fold could be attributed to either the lower effectiveness of some of these antibiotics, which was masked by the small volume of the microdilution method, the low sensitivity of the broth microdilution method or the increased total number of live bacterial cells. However, the CSLI guidelines have the same MIC breakpoints for both broth methods. This may require further studies using a larger number of bacterial strains to re-define the MIC breakpoints using both methods.

**Table 1.** MIC values of the broth micro- (I) and macrodilution (A) methods with (R) and without resazurin. The results are representative of four replicates in two independent assays. Values in bold are the more frequent values (n=3).

| Strains | Vancomycin MIC (mg/L) |  |  |  | Daptomycin MIC (mg/L) |  |  |  | Dalbavancin MIC (mg/L) |  |  |  |
| --- | --- | --- | --- | --- | --- | --- | --- | --- | --- | --- | --- | --- |
|  | I | I_R | A | A_R | I | I_R | A | A_R | I | I_R | A | A_R |
| MSSA (ATCC 25923) | 2 | 2 | 2 | 2 | **0.5-1** | **0.5-1** | 1 | 1 | 0.06 | 0.06 | 0.25 | 0.25 |
| MSSA (ATCC 29213) | 1 | 1 | **1-2** | 2 | 0.5-1 | **0.5-1** | **0.5-1** | 1 | **0.06-0.12** | **0.06-0.12** | 0.25 | 0.25 |
| MRSA (ATCC 43300) | 1 | 1 | **2-1** | 2 | **0.25-0.5** | 0.25-**0.5** | 0.5-**1** | 1 | 0.03-**0.06** | 0.03-**0.06** | 0.12-**0.25** | 0.25 |
| VISA (NARSA 17) | 8 | 8 | 8 | 8 | 4 | 4 | 4 | 8 | 0.25 | 0.25-**0.5** | 1 | 1 |
| hVISA (ATCC 700698) | 2 | 2 | 4 | 4 | 1 | **1-2** | 2 | 4 | 0.12 | 0.12-0.25 | 0.5 | 0.5 |
| VRSA (NARSA VRS3b) | 16 | **32-64** | >64 | >64 | 0.5 | 0.5 | **0.5-1** | 1 | 0.5-**1** | 1-2 | 2 | 4 |
| VRSA (NARSA VRS4) | >64 | >64 | >64 | >64 | 1 | 1 | 1 | 1 | 2 | 4 | 4 | 8 |

### 2. Evaluation of MIC by FACS

In order to assess the potency of these three antibiotics against the tested strains, replicates of the broth macrodilution MIC were analysed by FACS to detect and sort single cells. Single-cell sorting was performed to assess the actual percentage of live and dead cells using PI^43^ at 24 h endpoint. PI is a cell-impermeant, non-fluorescent dye that is only fluorescent when bound to bacterial DNA due to membrane damage. Positive controls of live and dead cells were used to identify the gates.

At each antibiotic concentration, the percentage of live cells for each strain was plotted and the results were compared to the resazurin macrodilution MIC values. An illustration of live and dead plots of dalbavancin treated MRSA is shown in **Figure S2**. At the MIC concentration, as determined by the broth macrodilution method, vancomycin MICs showed a range of 1.9–10.2% live cells based on the tested strains (Figure 1). However, daptomycin and dalbavancin showed a range of 17.7–62.9% and 7.5–77.6% live cells based on the strain (Figure 1), respectively. Consequently, the comparison between the visually detected MIC by broth macrodilution method and cell sorting of live/dead cells showed that the minimum inhibitory concentration represented from 22–98% of dead cells. This percentage varied based on the antibiotic and the tested strain, which cannot be identified using current MIC methods.

Interestingly, the vancomycin-treated hVISA strain evidenced to be a heterogeneous strain with a higher area under the curve than the MSSA and MRSA strains in concordance with the modified population analysis profile (PAP) method.^38,44^ The later requires extensive culture methods to detect vancomycin-associated bacterial heterogeneity, which has been proposed to be about ≤10^−5^ − 10^−6^ of the population, ^38,45^ this represents 0.001–0.0001% of bacterial cells in MIC assays. However, vancomycin showed 9.8% live persister cells with hVISA, and a plateau in the survival kinetics of 2–18.7% live persister cells with all tested strains, which is a higher percentage than previously predicted. A plateau in the survival of some bacterial cells was also noticed with daptomycin and dalbavancin with 5.4–17.4%, considering only VRSA strains, and 3.6–11.7%, considering all strains except VRSA, live persisters (Figure 1), respectively. Thus, persisters represented an average of 3.7±1.7% to 16±3.7%, which varied based on the strain and antibiotic.

**Figure 1.**
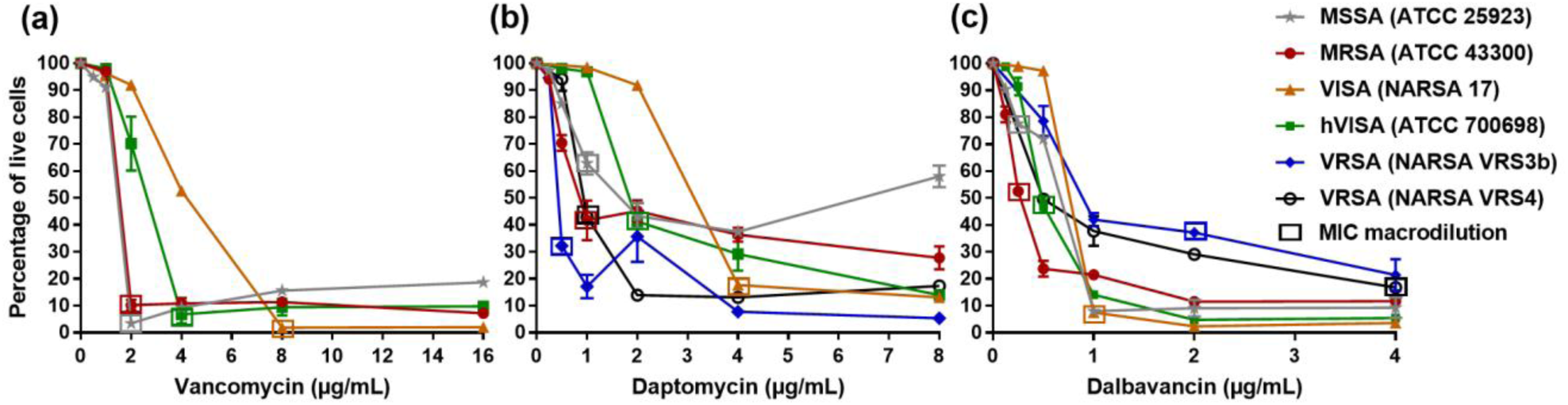
The percentage of different live bacterial strains as assessed by flow cytometry. Broth MIC macrodilution tubes were stained with PI and sorted for live/dead. The data were analysed based on the identified gates and the percentage of live cells were plotted. Square-labelled values are the broth MIC macrodilution concentrations. Data (n=3) are shown as means ± SD, some error bars are too small to be visible in the graph.

### 3. Detection of persisters and heterogeneity

The ability to scavenge and lower the generated ROS under stress was used to detect persisters^46^ and the results were compared to the detected live cells; therefore, replicates of the macrodilution broth method were stained with the membrane-permanent CELLROX^®^ dye and sorted. Control live and dead cells of all bacterial strains were stained and showed constant one population (**Figure S1d and Figure S1e**), a consensus gate was used as a reference ROS level (gate ‘ROS’) (**Figure S1c**). The ROS cell sorting results were analysed, compared to the ROS level of the strain and reference gate, the non-decreasing percentage of live cells, 3.7–16%, when treated at 2- to 16-fold the MIC and categorised in three groups: 1) the normal, 2) the heterogeneous and 3) the persistence-triggering group.

The normal ROS response showed a concentration-dependent increasing bacterial stress levels, which was shown with VRSA strains treated with dalbavancin (Figure 2a). At a sub-MIC concentration of dalbavancin, VRSA strains showed elevated ROS levels, which almost half of the population partially reduced at the MIC concentration. Above the MIC, the concentration-dependent potency of the antibiotic caused immediate death of most cells, preventing the management of the antibiotic-associated stress. However, a small percentage of cells (a sporadic population) maintained a low level of ROS (Figure 2a). Similarly, MRSA showed a higher stress reaction with dalbavancin than vancomycin at MIC concentrations, while VISA showed the opposite (Figure 2b).

Heterogeneous bacterial cells had a characteristic ROS response, where hVISA showed a very low ROS level to all the bacterial population when treated at sub-MIC concentration of vancomycin or dalbavancin and a slightly elevated stress level at the MIC concentration (Figure 2b). This ROS response was also identified with the VISA strain, a slow growing strain, which was shown in Figure 1 to have a higher area under curve than hVISA and MSSA which is confirmed to be a SCV strain (**Figure S3** and **Table S2**). Interestingly, MSSA showed a distinct persister population of low ROS level with a slightly different SSC when treated with vancomycin at 2-fold MIC, which is discussed in Figure 3 and **S4**. These results showed that heterogeneous bacterial cells had the ability to signal all the bacterial population to reduce their ROS level and avoid stress-associated cell damage.

Daptomycin, surprisingly, showed a persistent bacterial population with MSSA, MRSA and VRSA strains at 2-fold MIC (Figure 2c), and up to 16-fold MIC. Consistently, heterogeneous, slow growing and SCV strains, hVISA, VISA and MSSA, respectively, showed a low ROS level at sub-MIC concentrations of daptomycin, as previously identified with vancomycin and dalbavancin. This detected daptomycin-associated persistence may have partially contributed to the reported development of daptomycin resistance towards *S. aureus in vivo*.^47–49^ Interestingly, hVISA and MRSA showed different populations at sub-MIC and MIC concentrations of daptomycin and dalbavancin, respectively, (Figure 2c and 2b). These populations were of different low levels of ROS which may also suggest difference in scavenging mechanisms, which were the outcome of the potency of the antibiotics and the resistance of the strains.

It was noticed that some of the detected persister populations had a different SSC from normal cells (Figure 2). As the difference in SSC represents surface roughness, it was suspected that these populations may have a damaged cell membrane, which would affect the scattered fluorescence signal. Since they represent a very low percentage, bacterial cells were stressed by a quick 40% ethanol treatment followed by cell sorting and visualisation by microscopy. The cells were also stained with a fluorescent membrane dye (FM4-64) to prove that they had acquired surface granularity. The results showed that all cells had the same SSC when stained with PI or FM4-64 dye (**Figure S4a**), and only showed higher SSC when stained for ROS (**Figure S4b**). Microscopic imaging of the samples showed that most of the cells were not fluorescent (**Figure S4c**), as detected by FACS, and some cells showed a half-circled fluorescent ring, which probably had different light scattering than normal fluorescent cells. These results showed that the difference in scattering was probably due to the ROS localisation rather than a membrane damage.

**Figure 2.**
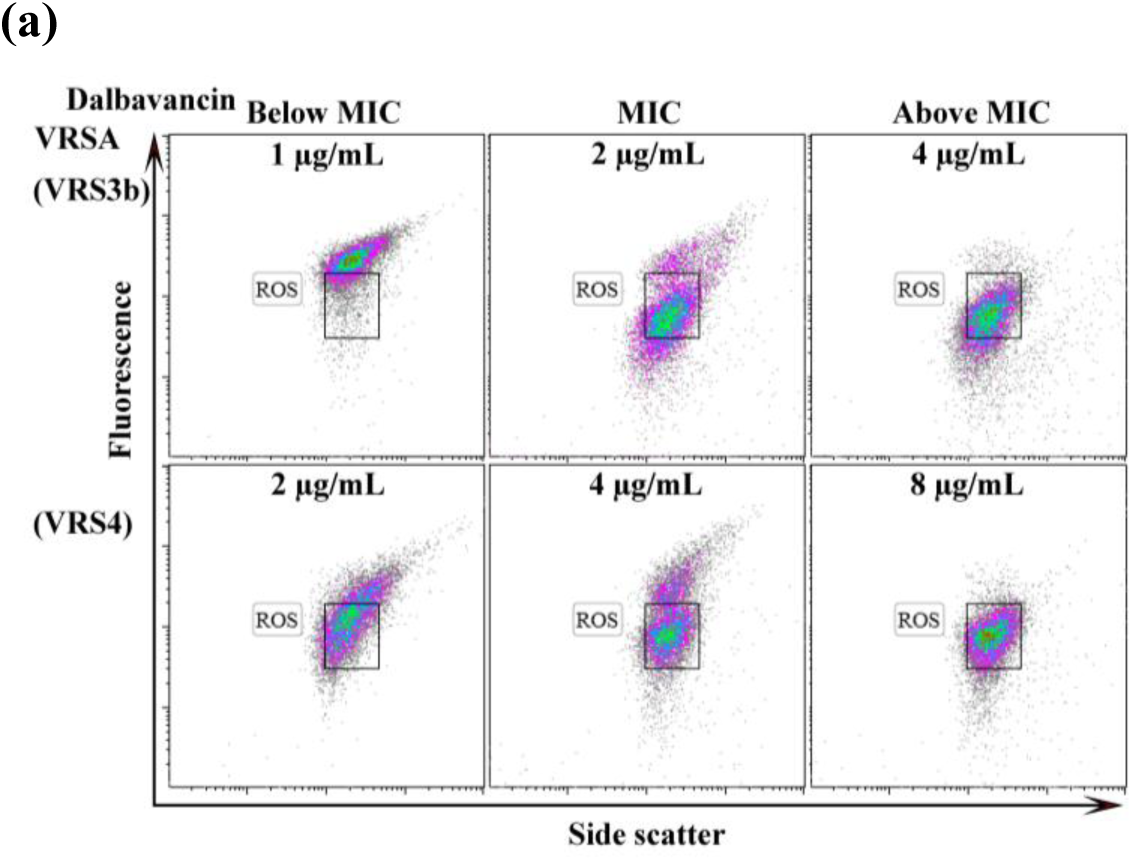
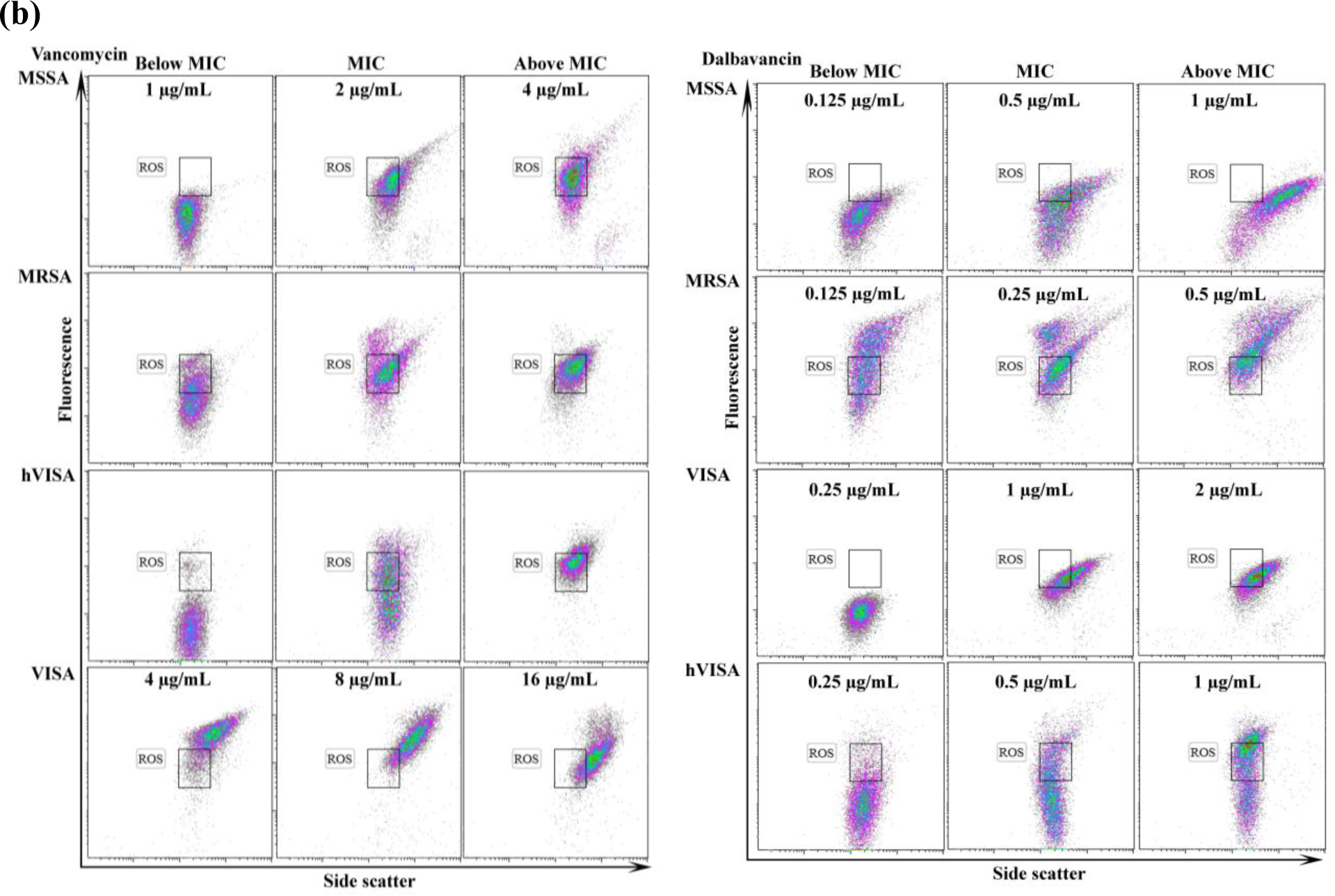
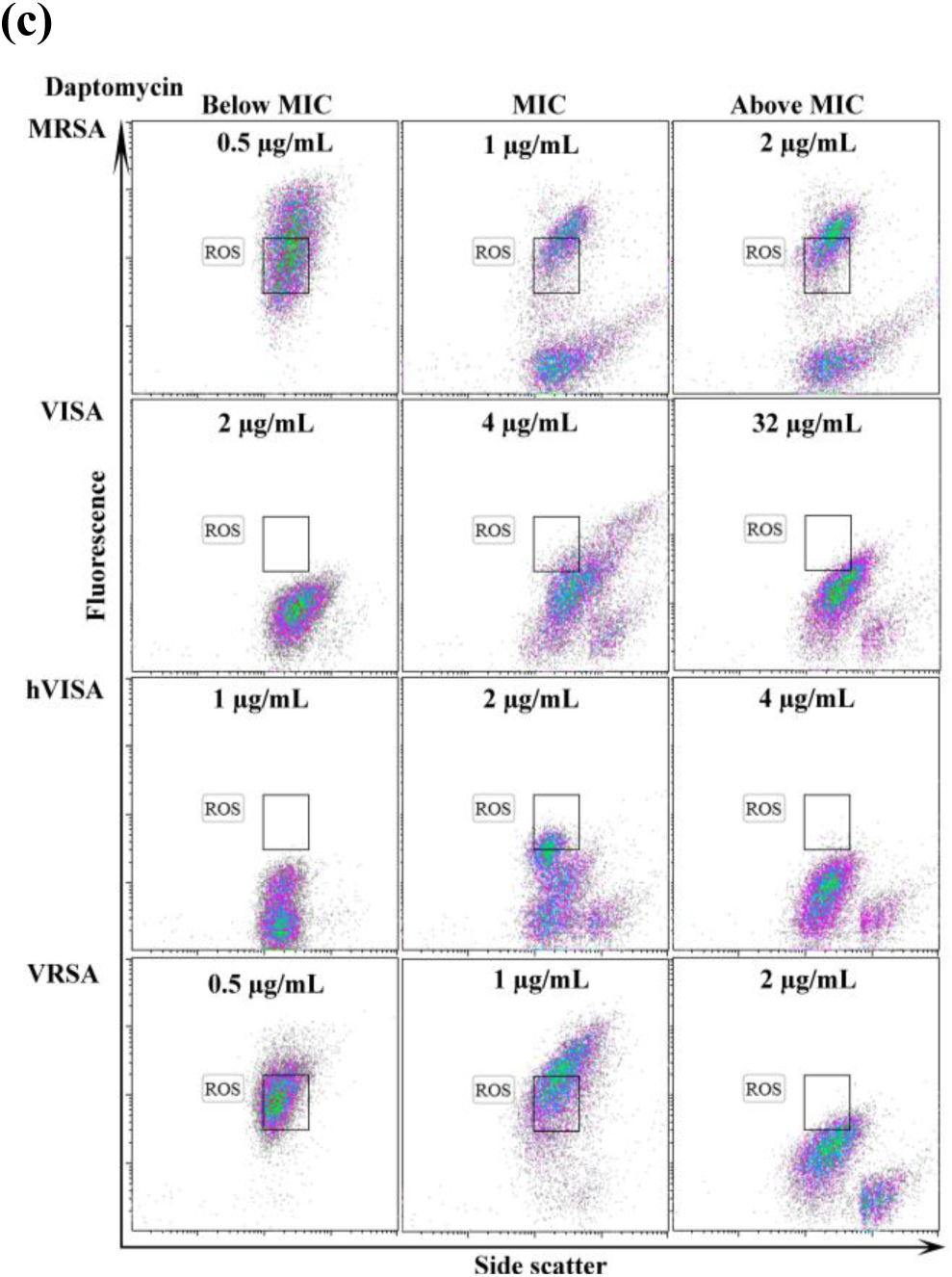
Cell sorting of the ROS generated under the treatment of antibiotics at different concentrations. ROS gate is the normal free radicals level assessed in the strains. **a)** A concentration-dependent increasing ROS response of bacterial cells. **b)** The ROS response of vancomycin and dalbavancin. **c)** The ROS response of daptomycin and its associated persisters. Data (n=2) are shown to be reproducible over two independent experiments.

To confirm the detection of SCV, a heterogeneous small colony with a parent colony and a parent colony only of MSSA were selected, treated with vancomycin and processed for detection (**Figure S3**). Cell sorting of the MSSA parent colony containing samples were previously demonstrated in Figure 2b, the SCV mixed colonies samples are shown in Figure 3. Samples were first stained with PI only and showed about 4% live population (gate R), representing 0.27% (gate P) on the SSC. Replicate samples were stained with ROS only and showed about 3.6% of the cells differentiated into higher surface roughness with low ROS levels (red cells, SSC, gate P) (Figure 3). PI was then added on the same ROS treated samples and proved that these persisters were live (gate L and R) (Figure 3). The SCV mixed sample showed slightly higher percentage of persisters when stained for ROS only compared to Figure 2b. This suggested that the increase in the SCV cells increased the persister population. It also confirmed that the 3.7–16% of live cells are persisters.

**Figure 3.**
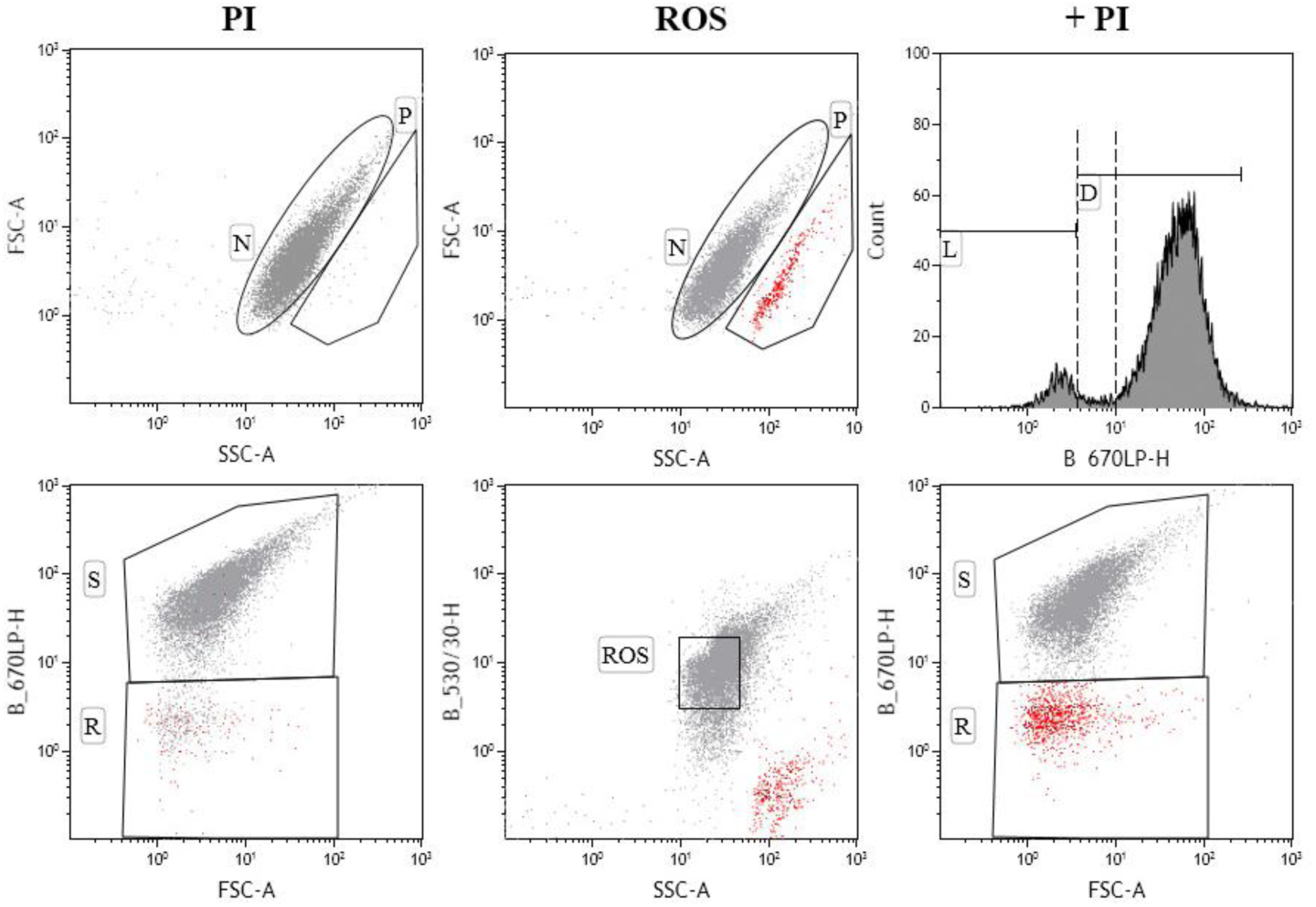
Detection of *S. aureus* SCV strain (ATCC 25923). The cells were treated with 2 mg/L vancomycin and stained with PI and CELLROX. The peak tail of gate D (in between the dashed lines) presents the partially damaged cells. Data (n=2) are shown to be reproducible over two independent experiments.

### 4. Re-evaluation of the efficacy of antibiotics

Even though MIC reflects only the bacterial inhibition, either in a static or cidal mode of action, but the inhibition of the growth was shown to not be consistent across tested antibiotics and strains. Therefore, the three antibiotics were re-evaluated for potency based on the death of at least 85% of the bacterial cells, with taking into consideration the formation of persisters. A heat map was plotted to visualise the difference in the potency of these antibiotics using MIC, live/death cell sorting and live/death with persistence detection (**Figure S5**). The MIC concentration of vancomycin showed ≥85% dead cells with all tested strains, and triggered a small persistent population with only MSSA. Whereas, daptomycin required a higher concentration than the MIC across almost all the strains and showed high levels of persisters. Daptomycin also failed to kill 85% of the MSSA and MRSA cells up to 16 mg/L (Table 2). Dalbavancin, however, showed more effective killing against VISA and VRSA (VRS4) strains (Table 2). Accordingly, the potency of these antibiotics varied from the culture-based MIC values, and triggered a concentration- or antibiotic-dependent persisters. This evidenced the limitations of the MIC methods^50^ to assess the efficacy of antibiotics, which can be re-evaluated based on both killing potency, rather than visual inhibition, and persistence formation to help reduce antimicrobial resistance.

**Table 2.** FACS-based evaluation of antibiotics based on live/dead percentage and persisters formation. Concentrations of the antibiotics presented showed ≥85% dead cells, values in bold are higher than MIC values detected by the resazurin macrodilution method, orange-highlighted values indicated the detection of persisters by ROS sorting.

| MIC FACS<br>($>85\%$ dead)<br>(mg/L) | MSSA<br>(ATCC<br>25923) | MRSA<br>(ATCC<br>43300) | VISA<br>(NARSA<br>17) | hVISA<br>(ATCC<br>700698) | VRSA<br>(NARSA<br>VRS3b) | VRSA<br>(NARSA<br>VRS4) |
| --- | --- | --- | --- | --- | --- | --- |
| Vancomycin | 2 | 2 | 8<sup>a</sup> | 4 | N/A | N/A |
| Daptomycin | $\geq 4$ (60%) | $\geq 1$ (60%) | 8 | **8** | **4** | **2** |
| Dalbavancin | **2** | **2** | 1<sup>b</sup> | **1<sup>c</sup>** | **16** | 8 |

<sup>a</sup> Represents 98% dead cells, <sup>b</sup> represents 90% dead cells and <sup>c</sup> the increase of the concentration to 2 mg/L had 95% dead cells and did not show a population of persistence.

## Conclusion

The current standardised MIC methods rely on visual detection, which does not reveal the biological state of the bacterial cells.^51^ In this study, testing of antibiotics with the broth macrodilution method was shown to be more sensitive, potentially due to the presence of a higher total number of cells. The use of resazurin increased the sensitivity of the broth macrodilution method, which was mainly dependent on the concentration of the dye. A thorough quantification of live, dead and persister cells was achieved with single-cell sorting, which demonstrated different efficacy of tested antibiotics and strains compared to MIC. Vancomycin showed to be very effective against MSSA, MRSA and VISA strains, while daptomycin failed to kill more than 60% of the population of MSSA and MRSA using cell sorting. Daptomycin also triggered the development of a persistent population with most of the tested strains, which may provide an explanation to the rapid development of resistance *in vivo*. The results also showed a higher percentage of persistence and heterogeneity in bacterial populations than is currently estimated. Therefore, assessment of the current standard MIC methods may be required to evaluate the efficacy of antibiotics. This also requires an extensive evaluation of a higher number of bacterial strains against all antibiotics to detect bacterial heterogeneity and guide for choice of the most suitable antibiotic treatment.

## Acknowledgements and funding

The authors acknowledge Prof. Mark Schembri for reviewing the manuscript. The authors also acknowledge the National Health and Medical Research Council (NHMRC) Project Grants APP631632 and APP1026922 for partially funding this work. M.M.H. is supported by the University of Queensland International PhD scholarship. M.A.C. is a NHMRC Principal Research Fellow (APP1059354) and currently holds a fractional Professorial Research Fellow appointment at the University of Queensland with his remaining time as CEO of Inflazome Ltd. The following reagent was provided by the Network on Antimicrobial Resistance in *Staphylococcus aureus* (NARSA) for distribution by BEI Resources, NIAID, NIH: NARSA STRAINS *Staphylococcus aureus*, Strain NARSA 17 (HIP06297, NR-45868), VRS3b (HIP13419, NR-46413) and VRS4 (HIP14300, NR-46414).

## Transparency declarations

None to declare.

